# Adaptable pulsatile flow generated by quantitative imaging of stem-cell derived cardiomyocytes for disease modeling

**DOI:** 10.1101/2020.03.20.000752

**Authors:** Tongcheng Qian, Daniel A. Gil, Emmanuel Contreras Guzman, Benjamin D. Gastfriend, Kelsey E. Tweed, Sean P. Palecek, Melissa C. Skala

**Author notes:** Correspondence should be addressed to T.Q or M.C.S.

## Abstract

Endothelial cells (EC) *in vivo* are continuously exposed to a mechanical microenvironment from blood flow, and fluidic shear stress plays an important role in EC behavior. New approaches to generate physiologically and pathologically relevant pulsatile flows are needed to understand EC behavior under different shear stress regimes. Here, we demonstrate an adaptable pump (Adapt-Pump) platform for generating pulsatile flows via quantitative imaging of human pluripotent stem cell-derived cardiac spheroids (CS). Pulsatile flows generated from the Adapt-Pump system can recapitulate unique CS contraction characteristics, accurately model responses to clinically relevant drugs, and simulate CS contraction changes in response to fluidic mechanical stimulation. We discovered that ECs differentiated under a long QT syndrome derived pathological pulsatile flow exhibit abnormal EC monolayer organization. This Adapt-Pump platform provides a powerful tool for modeling the cardiovascular system and improving our understanding of EC behavior under different mechanical microenvironments.

## Introduction

The microenvironment of blood vessels *in vivo* includes constant exposure to shear stress from blood flow, which plays an important role in vascular formation and in maintaining vascular function^1, 2^ Pulsatile flow is generated by the periodic contraction of the left ventricle and maintained throughout the arteries, as well as in selected areas of veins and capillaries^3^. Blood flow is tightly correlated to ventricular contractility^4^, so abnormal heart rhythm impacts peripheral blood flow pattern^5^. Human pluripotent stem cell (hPSC)-derived cardiomyocytes have been used to model the effects of genetic mutations on cardiomyocyte function, and are capable of recapitulating pathophysiological phenotypes^6^. However, effects of these mutations on the broader cardiovascular system, including blood vessels, are not captured in these hPSC-derived cardiomyocyte disease models. Endothelial cells (ECs) line the lumen of blood vessels, so pulsatile flow provides a more relevant *in vivo* microenvironment for these ECs compared to constant flow. Numerous studies have shown that pulsatile flow and constant flow have different effects on ECs *in vitro,* including EC morphology and alignment^7, 8^, gene expression^9, 10^, distribution of adhesion proteins^8^, cell proliferation, and apoptosis^11^. The pulsation frequency and peak amplitude of the pulsatile flow also plays an important role in EC morphology and protein expression^7, 8^. Additionally, fluidic shear stress significantly affects differentiation efficiency and related gene expression during the differentiation process^12, 13^, but the effects of pulsatile shear stress have not been well characterized. EC differentiation is involved in multiple blood vessel formation processes, including vasculogenesis and angiogenesis^14, 15^, and these new vascular formation processes are crucially affected by shear stress^16, 17^. Hence, it is important to develop cardiovascular models with more biologically relevant pulsatile flow profiles to better model vascular development and disease.

Previous methods to study EC behavior *in vitro* have generated pulsatile shear stress using pneumatic or piezoelectric pressure pumps and simulated waveforms^8, 18, 19^. However, the underlying pump technologies of these previous approaches lack the temporal resolution to recapitulate the frequency content of the time-dependent flow generated by heart contractions. Additionally, the simulated pulsatile waveforms of these previous studies cannot provide real-time changes in waveforms due to feedback flow or predict heart contraction in response to drug treatments for cardiovascular disease. Although previous studies have shown that pulsatile flows change the function and morphology of ECs compared to constant flows, the effects of variations in pulsatile flow, including pathological flows such as pulsatile flow from long QT syndrome (LQTS), on ECs are poorly understood. Hence, there is a need to generate adaptable pulsatile flows to study the role of variable pulsatile mechanical stimulation in vascular development, structure, function, and pathology.

Here, we demonstrate a proof-of-concept adaptable pump (Adapt-Pump) system to generate physiologically and pathologically relevant pulsatile flows based on the contractions of hPSC-derived cardiac spheroids (CSs). We developed quantitative imaging-based signal transduction to record the spontaneous periodic contractions of CSs and actuate the contraction waveforms with high fidelity as microfluidic pulsatile flow. We used Adapt-Pump to generate pulsatile flow from CSs derived from different hPSC lines, including healthy human embryonic stem cell (hESC)-derived CSs and patient-specific LQTS induced pluripotent stem cell (iPSC)-derived CSs. We then resolved the effects of drug treatment on CS contractions, and accurately recapitulated these contractions in microfluidic pulsatile flow. We also demonstrated real-time Adapt-Pump (rtAdapt-Pump) and resolved the instantaneous response of CS contraction to microfluidic mechanical stimulation. Adapt-Pump was also used to apply pulsatile flow to EC progenitors, and we discovered that LQTS-derived pulsatile flow induced abnormal cell organization during EC differentiation. This Adapt-Pump system provides a powerful new tool to generate physiologically or pathologically relevant pulsatile flow from hPSC-derived CSs. Adapt-Pump can be used to investigate integrated cardiomyocyte and EC behavior under various mechanical microenvironments resulting from cardiac disease in response to clinically-relevant drug treatments.

## Results

### Workflow to generate pulsatile flow from hPSC-derived cardiac spheroids for disease modeling

hPSCs can differentiate into specialized cell types to facilitate studying development, modeling disease, and screening drugs ^20, 21^. Here, we demonstrate a proof-of-concept Adapt-Pump system to generate pulsatile flow from hPSC-derived CSs. We showcase three applications with this Adapt-Pump system. First, we recapitulate pulsatile flow changes after clinically-relevant drug treatment to hPSC-derived CSs. Second, we demonstrate that the Adapt-Pump system can generate real-time pulsatile flow from CS contraction in response to feedback mechanical stimulation. Third, we perform EC differentiation under a pathological mechanical microenvironment derived from congenital LQTS iPSC-CSs. These applications demonstrate the use of the Adapt-Pump to generate pulsatile flows from different hPSC lines including patient-specific iPSCs, monitor real-time cardiac contraction in response to feedback fluidic flow, and perform studies of integrated cardiomyocyte-EC behavior across a range of fluidic mechanical microenvironments including those that mimic cardiac diseases.

The workflow is shown in Fig. 1. First, hPSCs were differentiated into cardiac progenitors using an established small-molecule-mediated differentiation protocol^22^. CSs were generated from a single-cell methylcellulose suspension of cardiac progenitors. After 48 hours, aggregated CSs were embedded into Matrigel and plated on ibidi coverslip dishes for imaging (Fig. 1a). Brightfield image time-series were acquired of the CS contractions, and quantitative image analysis was used to convert these images into a contraction waveform. Briefly, the 2D area (mm^2^) of the CS was quantified in each frame using image segmentation. The contraction waveform then was calculated as the change in area relative to the area of the fully relaxed CS (Fig. 1b). A scaling factor *C* was used to convert the contraction waveform into a pressure waveform in the range of 0 to 800 mbar. Finally, this pressure waveform was sent to a highspeed pump to generate pulsatile flow onto EC progenitors that lined a confluent monolayer in a medium-filled channel (Fig. 1c). Additional details regarding the experimental system are described in the Materials and Methods section.

**Figure 1.**
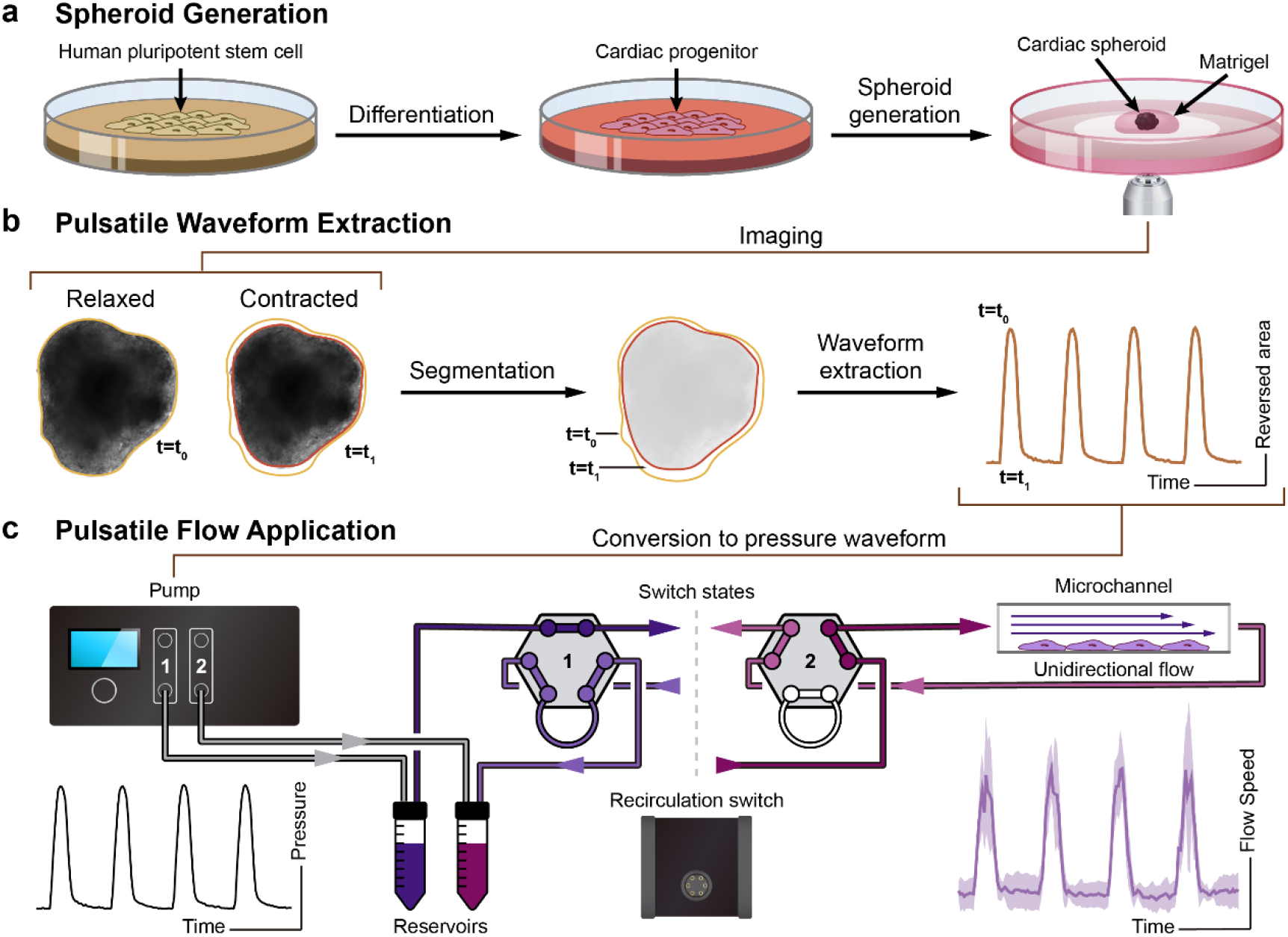
Schematic of workflow for generating pulsatile flows. (**a**) Singularized hPSCs are seeded and expanded for 2 days in mTeSR1. Differentiation to cardiac progenitors follows a previous protocol with a sequential Wnt activation and inhibition^22^. Cardiac spheroids (CS) are generated from cardiac progenitors and embedded in Matrigel for further cardiac contraction imaging. (**b**) hPSC-derived CSs are imaged. A custom script is used to analyze these time-series images and extract the pulsatile waveform from the CS contractions. (**c**) The contraction waveforms are converted to pulsatile waveforms and sent to the Elveflow OB1 pump. Elveflow MUX Injection system is used to recirculate cell culture medium. Fluidic pulsatile shear stress is applied to EC progenitors seeded in ibidi μ-slides for the EC differentiation study.

### Fluorescent bead streak analysis and shear stress simulation

The transit of fluorescence beads has been commonly used to characterize fluidic flow profiles and shear stress regimes at fluid-substrate boundaries^23^. Quantifying fluid speeds of over 1 cm/s exceeds the limitations of commonly used fluorescence microscopy equipment and standard single-particle tracking approaches. To fully characterize the high-speed (>1 cm/s) pulsatile flow profiles, we developed a method based on particle streak velocimetry to measure the instantaneous fluid speed using epifluorescence microscopy ^24^. The particle streak was tracked by using 4 μm fluorescent microbeads in phenol red-free medium in a 0.8 mm-ibidi μ-slide under pressures ranging from 50 mbar to 200 mbar. Time-series images of bead flow were recorded with focal point of the objective at z-positions of 0.5 mm, 0.6 mm, 0.7 mm, and 0.8 mm (z-position 0.0 mm is the bottom surface of the channel) at 200 frames per second (fps) (Fig. 2a, supplemental videos 1-4). A custom MATLAB script analyzed the image time-series and calculated fluorescent bead speeds. Briefly, an intensity threshold on raw images (Fig. 2b-raw) removed out-of-focus streaks, small objects, and streaks touching the image border. These thresholds were also used to select beads near the focal position, which were sharper and brighter than those farther away. Bead speed were then calculated from the length of each processed streak divided by the frame period (5 ms) (Fig. 2b-processed). The parabolic velocity profile of laminar flow in a closed rectangular channel has been characterized in previous studies^25^. A quadratic function was performed to confirm the parabolic relationship between bead speed and axial position in the 0.8 mm ibidi μ-slide. The parabolic fits for speed profiles and R^2^ values for 50 to 200 mbar input pressures are above 0.95 as shown in Fig. 2c. Input pressure of 800 mbar yields a shear stress of approximately 11 dyne/cm^2^ in the ibidi 0.8 mm μ-slide (Fig. 2c-e, Fig. S1). These characterization studies confirm that varying levels of laminar flow can be generated by the Adapt-Pump system.

**Figure 2.**
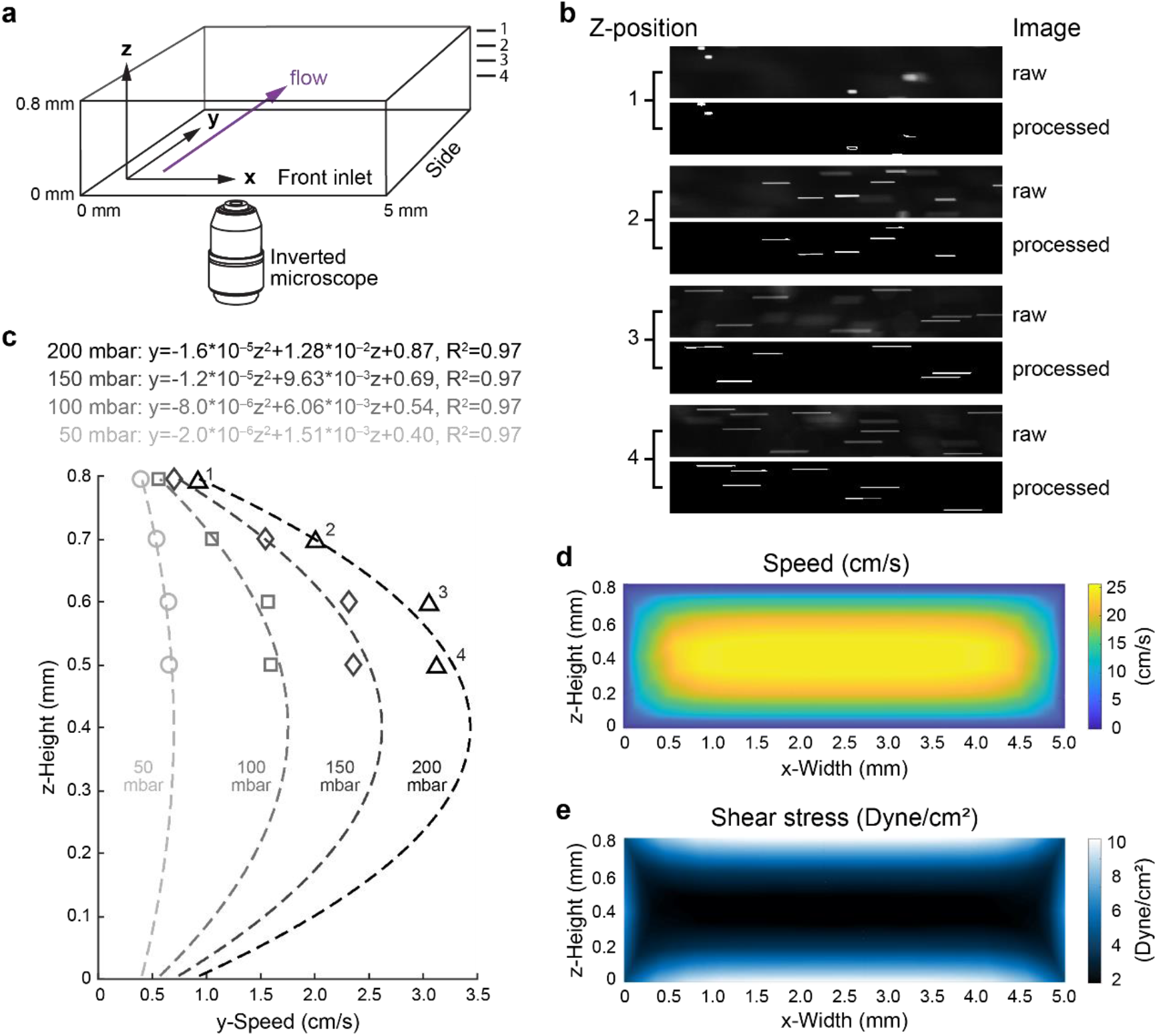
Fluorescent bead speed measurement and shear stress simulation. (**a**) Schematic of ibidi μ-slide channel used for flowing bead streak measurement and shear stress simulation. Dimensions as labelled. Flow direction is indicated in purple arrow. Z-positions are 0.8 mm, 0.7 mm, 0.6 mm, and 0.5 mm for measurements labeled “1”, “2”, “3”, “4”. (**b**) Representative images of fluorescent bead streaks (raw and after image processing steps, or “processed”) from the z-positions 0.8 mm (“1”), 0.7 mm (“2”), 0.6 mm (“3”), and 0.5 mm (“4”) generated at 200 mbar pressure. (**c**) A custom algorithm was used to calculate the fluorescence bead speed at different channel z-positions (1-4). Pressures are indicated. Simulated flow speed (**d**) and shear stress (**e**) at 800 mbar output from Elveflow pump are presented in front view, respectively.

### Cardiac spheroid contraction waveform and pulsatile flow characterization

As described in Fig. 1, pulsatile flows were generated from CSs differentiated from multiple hPSC lines, including H9 hESCs, H1 hESCs, and LQTS iPSC UCSD102i-2-1 (LQTS iPSC). A representative brightfield image of a CS (Fig. 3a), an image time-series of CS contraction (Supplemental video 5), and CS contraction waveforms for each hPSC line (Fig. 3b) showed unique contraction patterns between normal and pathological conditions. Normal ESC-CSs (differentiated from H9 and H1 hESCs) exhibited uniform contractions, but the LQTS-CSs exhibited periodic elongated contractions (indicated by the red arrow in Fig. 3b, supplemental videos 5-7) followed by a series of fast contractions (outlined by the red rectangle in Fig. 3b). This irregular contraction pattern is consistent with torsades de pointes (TdP), the clinically observed abnormal heart rhythm characteristic of LQTS^26^. Normalized CS contraction amplitudes of H1-CS and H9-CS were different from that of LQTS-CS. Contraction frequencies for CSs from all three hPSC lines were different from each other. The H9-CS contraction duration was significantly shorter than those of H1-CS LQTS-CS. Even though LQTS-CS exhibited higher contraction frequency, LQTS-CS had longer average contraction duration than H9-CS (Fig. S2). The longer contraction duration for LQTS-CS is attributed to the periodic long contractions. These contraction waveforms were then used to generate pulsatile microfluidic flow that fully recapitulates the corresponding contraction waveforms at high fidelity (Fig. 3c, d, Supplemental Videos 8-10) with Pearson’s r values above 0.75 (Supplemental Table 1).

**Figure 3.**
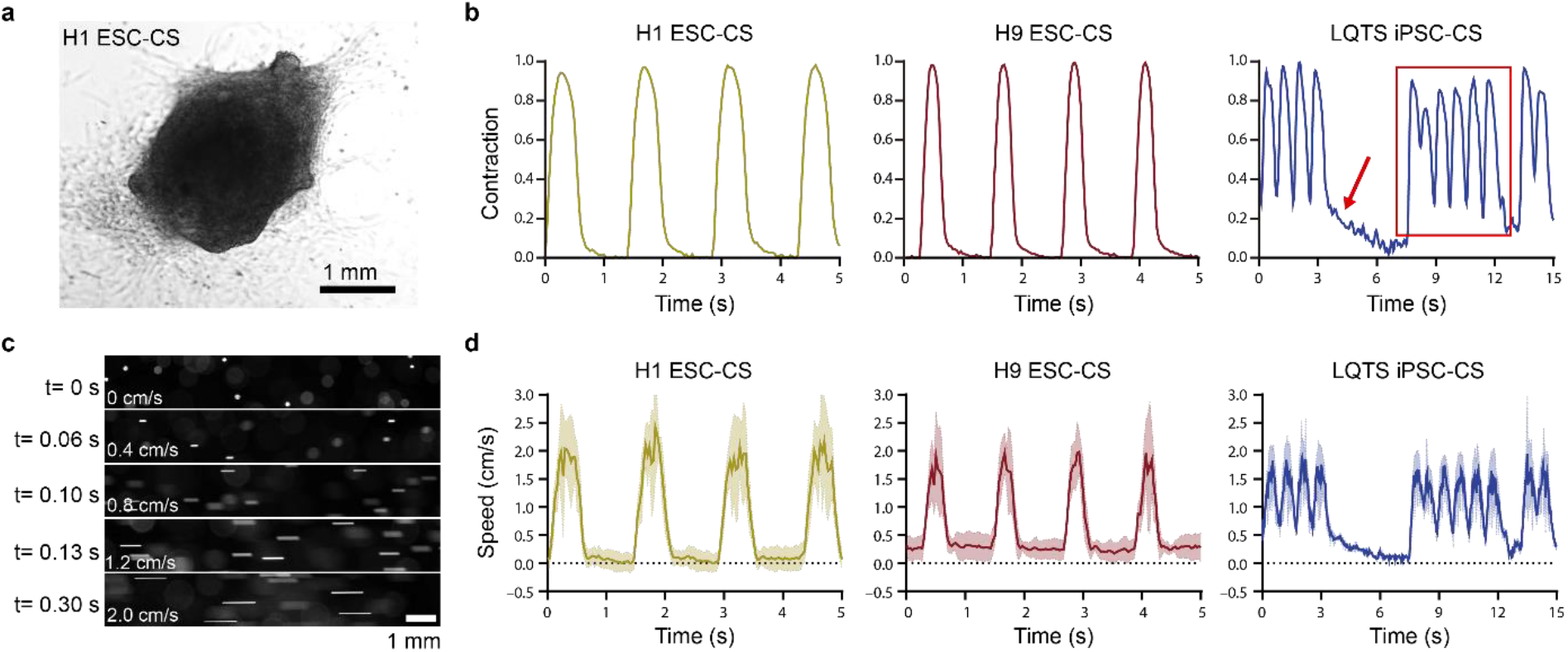
Characterization of hPSC-CS contraction waveforms and fluidic pulsatile waveforms. (**a**) Representative brightfield image of H1 ESC-derived CS. (**b**) Representative hPSC-derived CS contraction waveforms of H1 ESC-CSs, H9 ESC-CSs, and LTQS iPSC-CSs were extracted by a custom algorithm. Peak contraction magnitude was normalized to 1. Red arrow indicates long duration contraction and red rectangle indicates a series of fast contractions. (**c**) 150 mbar pressure was sent from an Elveflow pump as the normalized contraction magnitude 1. Representative fluorescence microscopy images of flowing beads at different time points. (**d**) A custom algorithm was used to calculate the speed of flowing beads as driven by the pulsatile contraction waveforms of H1 ESC-CSs, H9 ESC-CSs, and LQTS iPSC-CSs from fluorescence microscopy. Waveforms include mean (solid line) and SEM (shaded line).

Pulsatile flows were also generated based on the response of periodic contraction of hPSC-CSs to drug treatment (Fig. 4). Verapamil^27^ and nifedipine^28^ are calcium antagonists used to treat angina^29^, hypertension^29^, and LQTS^30^. 10 minutes of 1 μM verapamil treatment decreased contraction amplitude by over 20%, decreased contraction frequency from 0.9 Hz to below 0.7 Hz, and increased contraction duration by over 20% in H9-CSs (Fig. 4a, b, supplemental videos 11-12). 10 minutes of 1 μM nifedipine treatment significantly dropped the contraction amplitude and decreased contraction frequency from above 3 Hz to below 2 Hz, and increased contraction duration by nearly 100% in LQTS-CSs (Fig. 4d, e, supplemental videos 13-14). This is consistent with a previous study showing that nifedipine can inhibit LQTS-triggered arrhythmias in patients^31^. The Adapt-Pump was then used to generate pulsatile microfluidic flow that reflects the response of CS to drug treatment (Fig. 4c, f, Table 1, supplemental videos 15-18). Therefore, this system could be used for to resolve the effects on EC behavior under fluidic mechanical environment caused by abnormal cardiac contractility due to cardiovascular disease or drug treatment.

**Figure 4.**
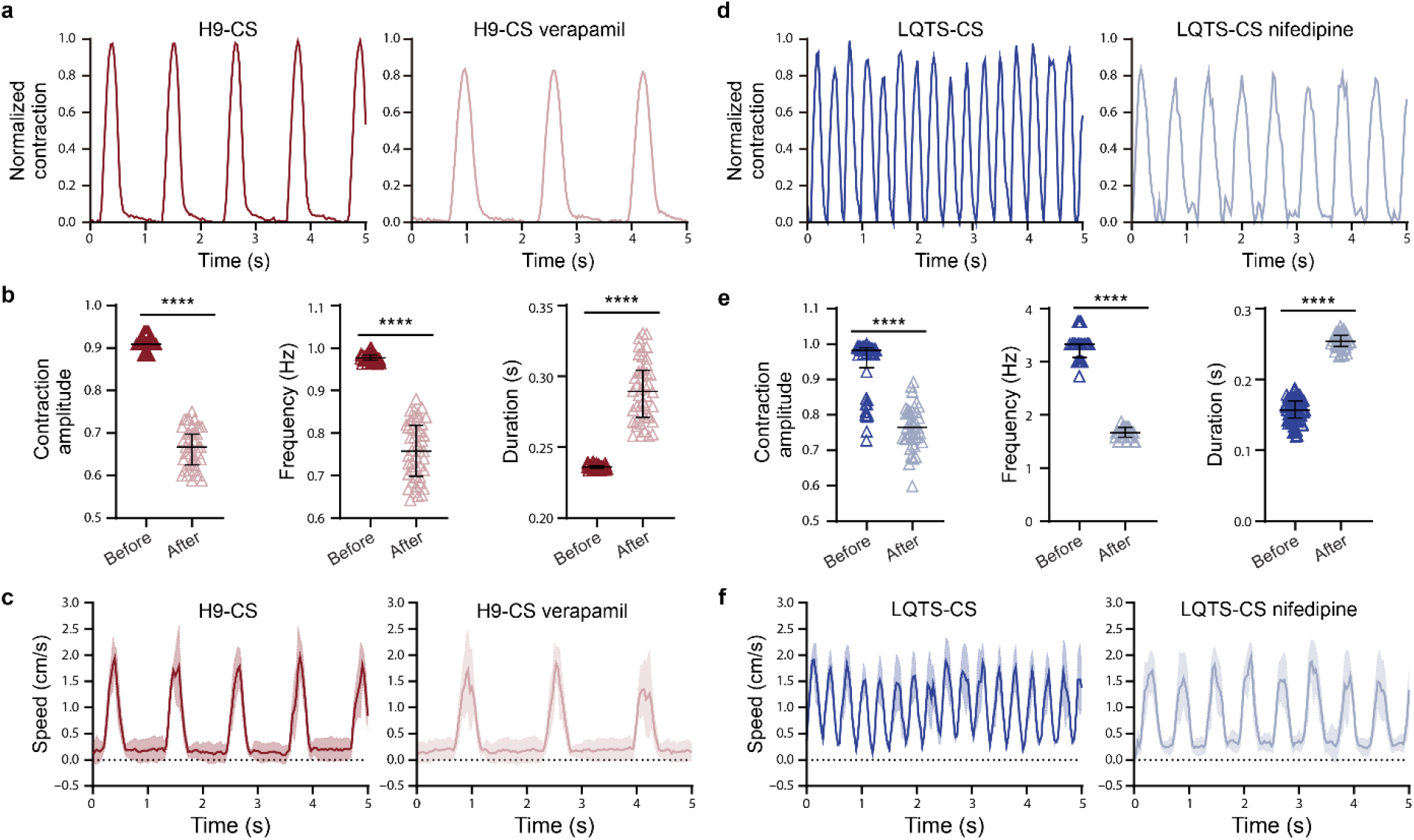
Characterization of hPSC-CS contraction waveforms and fluidic pulsatile waveforms before and after drug treatment. (**a**) Representative H9 hESC-CS contractions before and after 10 minutes treatment of 1 μM verapamil were recorded with a real-time microscopy and calculated by a custom algorithm. (**b**) Normalized contraction amplitude, contraction frequency, and contraction duration of H9 ESC-CSs before and after verapamil treatment (black lines indicate mean ± SEM). The contraction waveforms were calculated from individual contractions over 60 seconds. (**c**) 150 mbar pressure was sent from an Elveflow pump as the normalized contraction magnitude 1. Representative bead speeds were recorded with a real-time microscopy and calculated by a custom algorithm. (**d**) Representative LQTS iPSC-CS contractions before and after 10 minutes treatment with 1 μM nifedipine were recorded with real-time microscopy and calculated by a custom algorithm. (**e**) Normalized contraction amplitude, contraction frequency, and contraction duration of LQTS iPSC-CS before and after nifedipine treatment were presented as mean ± SEM. The contraction waveforms were calculated from individual contractions over 60 seconds. (**f**) Representative bead speeds. Statistical significance was determined by Student’s t test (two-tailed) between two groups. **** *P* < 0.0001.

### Real-time CS contraction feedback in response to fluidic mechanical stimulation

In addition to generating pulsatile flows from different hPSC-CSs and response to drug treatment, we also used the Adapt-Pump system to demonstrate real-time CS contraction feedback to fluid flow. CS contractions were monitored while the CSs were exposed to the fluid flow generated from their own contractions. Our proof-of-concept demonstration mimics the real time cardiac contraction response to fluidic mechanical stimulation. As shown in Fig. 5a, the rtAdapt-Pump is an organ-on-a-chip system that includes a CS embedded in Matrigel underneath a microfluidic channel that circulates cell culture medium, a live cell imaging system, a real-time image processing system, and a microfluidic pump. For the static condition, CS contractions were imaged over 30 seconds to record the contraction waveforms before flow was applied (Fig. 5b black line). For the flow condition, CS contraction were monitored over 30 seconds in the presence of fluidic flow based on a contraction waveform extracted in real time from the same CS (Fig. 5b, purple line). The contraction peak amplitude, contraction duration, and area under the curve (AUC) of contractions in Fig. 5b are plotted in Fig. 5c. All three parameters in Fig. 5c were significantly higher under the flow condition compared to the static condition, which is consistent with a previous study that showed mechanical stimulation modulate cardiomyocyte contraction^32^. This rtAdapt-Pump system provides a cardiovascular system model that could enable studies to better understand integrated cardiomyocyte-EC behavior under a variety of feedback conditions.

**Figure 5.**
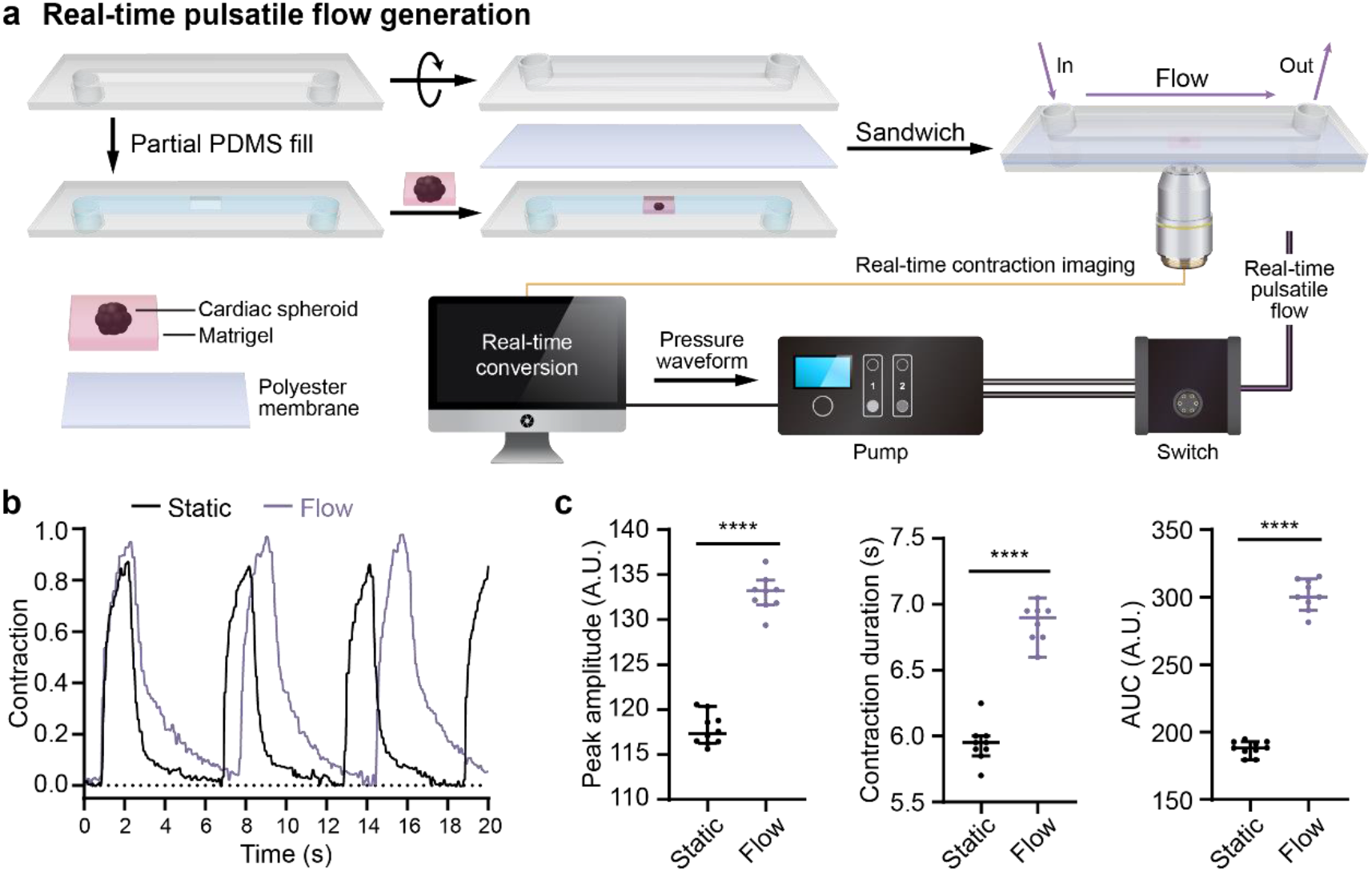
hPSC-CS contraction in response to fluidic mechanical stimulation. (**a**) Schematic of heart-on-a-chip modeling CS contraction feedback in reponse to fluidic mechanic stimulation. CSs were embedded in Matrigel and plated into the microchannel of a 0.8 mm ibidi sticky μ-slide (lower chamber). The lower chamber was sealed with a polyester membrane and covered with another 0.8 mm ibidi sticky μ-slide (upper chamber). The upper chamber was filled with CS culture medium. For contractions under static condition, CSs were first imaged with brightfield microscopy before applying fluidic flow to the spheroids. For contraction response upon to fluidic mechanic stimulation, CSs were imaged and the contraction waveforms were analyzed simultaneously by a computer. The pulsatile flow signal was instantaneously sent to the pump and the fluidic flow was applied to the CSs. The spheroid contractions in response to pulsatile flow were recorded by brightfield microscopy. All signals were analyzed by a custom algorithm. (**b**) Representative spheroid contraction waveforms before and after fluidic flow stimulation. (**c**) The peak contraction amplitude, contraction duration, and area under curve (AUC) for the contractions are presented as means ± SEM. All the contraction waveforms were calculated from the individual contractions over 30 seconds. Statistical significance was determined by Student’s t test (two-tailed) between two groups. **** *P* < 0.0001.

### Endothelial cells differentiated under LQTS pulsatile flow exhibit abnormal structures

Lastly, we also investigated pulsatile flow effects on hPSC differentiation to ECs. EC differentiation was monitored under physiological (H9-CS waveform) fluid flow or static conditions. ECs *in vivo* experience different pulsatile shear stress at different locations; for example, the shear stress in small arteries is over 12 dyne/cm^2^^33^. Therefore, an H9-CS pulsatile waveform with an average shear stress of 13.7 dyne/cm^2^ (Fig. S3a) was applied to EC progenitors at day 6 of differentiation. At this stage, CD34+ EC progenitors are multipotent and can differentiate into CD31+ ECs and smooth muscle cells^34^. After 48 hours further differentiation, ECs under shear stress had significantly higher differentiation efficiency, with approximately 85% CD31+ cells compared to 65% CD31+ cells under static conditions (Fig. 6a, b, red arrows indicate CD31 negative cells). CD31 expression levels were measured by flow cytometry without membrane permeabilization to reveal that ECs differentiated under static conditions had an approximately 30% higher mean fluorescence intensity of CD31 immunostaining compared to ECs differentiated under shear stress (Fig. 6c). However, CD31 mean fluorescence intensities for both static and shear differentiation conditions were similar after membrane permeabilization (Fig. 6d). The differences in labelled CD31 fluorescence levels are likely due to the subcellular distribution of CD31 (Fig. 6a). CD31 was largely localized to cell-cell junctions under static conditions, while more punctate CD31 was detectable in the cytoplasm of cells differentiated under shear stress (Fig. 6a). Pulsatile shear flow also increased the forward scatter of ECs by over 50%, which indicates larger ECs (Fig. 6e). These observations indicate that pulsatile flow can redistribute CD31 and alter the cell size, which is consistent with previous findings that pulsatile flow regulates EC protein localization^35^ and changes EC size by rearrangement of cellular components ^36^.

**Figure 6.**
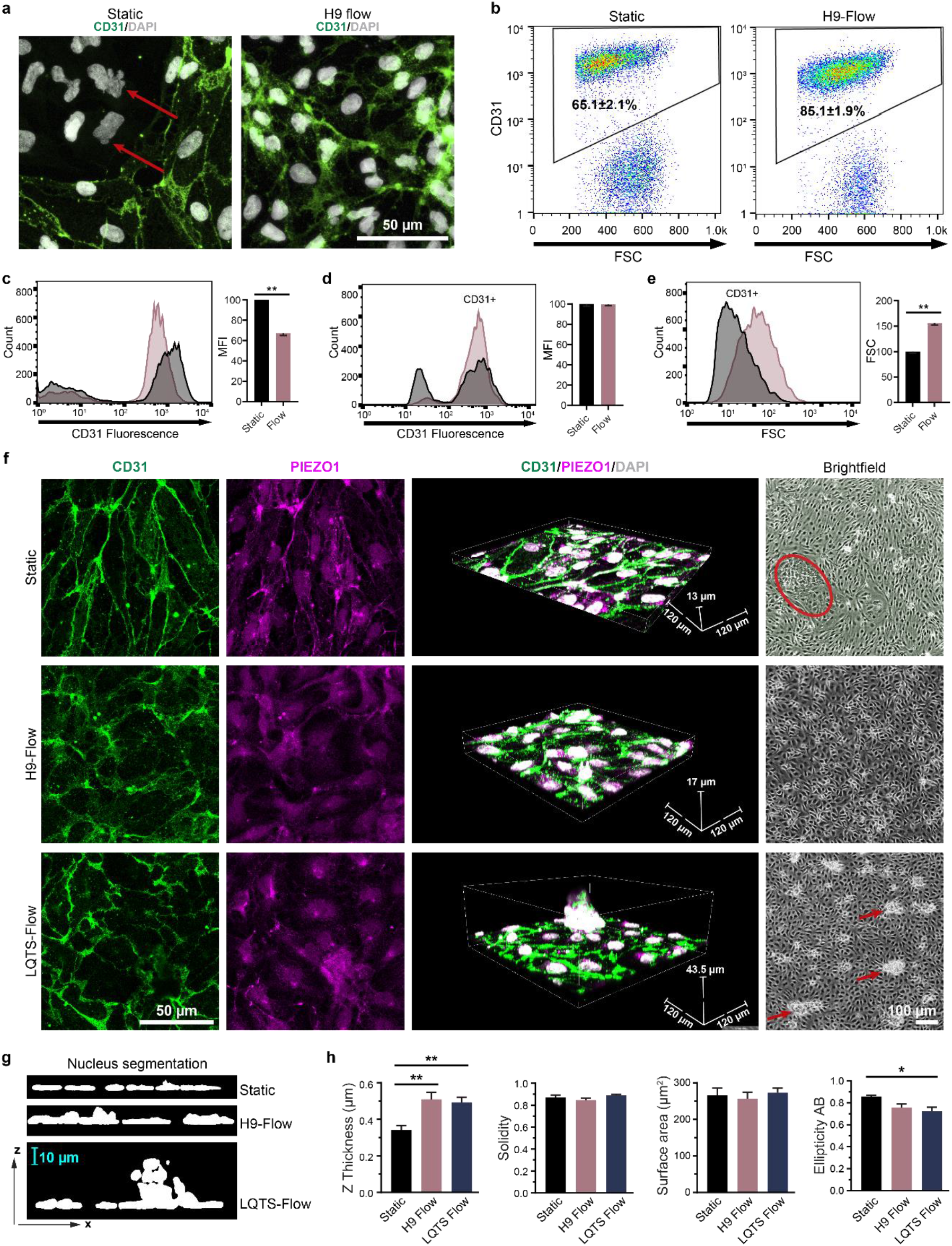
Fluidic pulsatile flow affects EC differentiation efficiency and morphology. (**a**) EC progenitors were differentiated under static conditions or under hPSC-derived pulsatile flow conditions as shown in Fig. 1c. After 48 hours of differentiation, ECs were immunofluorescently labelled with anti-CD31 (green) and stained with DAPI (white). Red arrows indicate non-endothelial cells. (**b**) EC populations were quantified by flow cytometry with CD31 labelling from three independent biological replicates and presented as means ± SEM. Extracellular CD31 expression level was quantified by flow cytometry without membrane permeabilization as the mean fluorescence intensity (MFI) of the CD31+ population. Statistical significance was determined by Student’s t test (two-tailed) (**c**). Total CD31 expression (MFI) (**d**) and EC size (forward scatter, FSC) (**e**) were quantified by flow cytometry with membrane permeabilization. The CD31+ population was gated and is shown in the histogram plot. Statistical significance was determined by Student’s t test (two-tailed) (**f**) ECs after differentiation under static conditions, H9-CS-derived pulsatile flow conditions, and LQTS-CS-derived pulsatile flow conditions. After 48 hours of differentiation, ECs were immunofluorescently labelled with anti-CD31 (green) and anti-PIEZO1 (magenta), and stained with DAPI (white). Labelled cells were imaged with a 0.5 μm Z step. Red circle indicates non-endothelial cell population and red arrows indicate abnormal EC clusters. (**g, h**) Images were reconstructed in ImageJ and nuclear morphology was quantified. Data were collected from a minimum of three independent differentiations and plotted as mean ± SEM. Statistical significance was determined by one-way analysis of variance (ANOVA) followed by Tukey’s post hoc tests. \**p*< 0.05, \*\**p*< 0.01.

Next, Adapt-Pump was used to determine whether pulsatile flows derived from normal ESC-CSs and LQTS iPSC-CSs can affect EC morphology or function. The average shear stress for both conditions was 13.7 dyne/cm^2^-(Fig. S3a). Flow was applied to EC progenitors at day 6 postdifferentiation for 48 hours, and ECs were assessed by immunofluorescence for CD31 and the mechanical sensor PIEZO1, and nuclear stain DAPI (Fig. 6f, supplemental videos 19-21). Cells differentiated under pulsatile flow (normal and LQTS) had more punctate CD31 and PIEZO1 with thicker nuclei compared to cells differentiated under static conditions (Fig. 6g, h). Morphology of EC nuclei differentiated under different conditions were quantified with labelled images from confocal microscopy. Nuclear thickness significantly increased under flow conditions. Nuclear ellipticity was also deceased under flow. However, there were no significant difference in solidity and surface area (Fig. 6h). ECs differentiated under LQTS pulsatile flow had randomly distributed CD31+ cell clusters on top of an endothelial cell monolayer (Fig. 6f bright field red arrow, g). Conversely, ECs differentiated under the static and H9-CS pulsatile flow conditions had uniform monolayers, especially ECs differentiated under H9-CS pulsatile flow (Fig. 6f bright field. Red circle indicates non-ECs under static condition). These experiments show that variations in pulsatile shear stress affects EC differentiation efficiency and EC monolayer organization, and demonstrate that the Adapt-Pump can be used for future studies of EC differentiation under a variety of fluidic mechanical microenvironments caused by different cardiac diseases.

## Discussion

Here, quantitative imaging of hPSC-CSs contraction profiles and a high-speed microfluidic pump were combined into the Adapt-Pump system to generate biologically-driven pulsatile flows to facilitate modeling the cardiovascular system. Previous methods to generate pulsatile flows were based on slow pressure pumps and simulated waveforms ^7, 35^, which could not fully replicate the *in vivo* mechanical microenvironment experienced by ECs. The Adapt-Pump system demonstrated here can recapitulate differences between normal and diseased cardiac waveforms including variable pulsatile magnitude, duration, and frequency at high temporal resolution. The Adapt-Pump system also captures dynamic changes in pulsatile flow after cardiac drug treatment that recapitulates *in vivo* cardiac drug response, and can be performed in real-time to understand feedback mechanisms in CS contraction. Overall, this system could be used to model the realtime response of heart contraction to drug treatment and feedback flow *in vitro,* investigate cardiomyocyte and EC behavior under mechanical microenvironments related to cardiac disease, and integrate into an organs-on-a-chip system for a biologically-driven circulatory system.

Genetically deficient iPSCs have been used for *in vitro* disease modeling, including models of LQTS^6^. We demonstrated that 3D CSs generated from different hPSCs recapitulate the heart rhythm characteristics of their genotype, including normal heart contraction from ESC-derived CSs and periodic elongated contraction from LQTS iPSC-derived CSs. Abnormal heart rhythm, resulting in abnormal flow, may affect EC biology^37^, but existing technologies to generate *in vitro* pulsatile flow lack biological relevance because they use synthetic heart contraction waveforms. Our method combines hPSC-CSs, high-speed quantitative image analysis, and fast signal transduction to generate pulsatile flow that adapts to the hPSC genotype and effects of drug treatment.

ECs *in vivo* are continuously exposed to fluidic mechanical stimulation from pulsatile blood flow^38^, and fluidic shear stress influences EC behavior and differentiation^12^. We discovered that pulsatile flow significantly enhances EC differentiation efficiency, which is consistent with a prior report that shear stress can induce EC differentiation^12^. We also found that pulsatile flow increases the thickness of nuclei, consistent with previous reports of changes in EC nuclear shape with fluid flow^39^. These changes in nuclear morphology could be an indication of the response of ECs to shear stress. Pulsatile flow also increased cell size and changed the distribution of the endothelial cell adhesion molecule CD31 and mechanical sensor PIEZO1, suggesting that EC structure is affected by pulsatile flow. Pulsation, including frequency and amplitude, plays an important role in EC protein expression and morphology^7, 8^. Disturbed flow is known to affect EC behavior, including EC turnover, low density lipoprotein permeability, atherogenesis, and gene expression^40–43^. However, the relationship between abnormal pulsatile flow and EC behavior has not been previously characterized due to the lack of appropriate methods to generate pulsatile flow that are relevant to cardiovascular disease. With the Adapt-Pump system presented here, pulsatile flow from LQTS was generated and applied to EC progenitors during differentiation, resulting in abnormal EC organization. However, LQTS patients do not continuously experience abnormal pulsatile flows over 48 hours, rather, these patients experience arrhythmias periodically throughout their lives. Long-term effects on EC biology from these abnormal pulsatile flows caused by congenital heart diseases is unclear, but our proof-of-concept EC differentiation studies show that abnormal pulsatile shear stress can dramatically affect EC behavior.

The hPSC-CS-derived pulsatile flows used in these studies also recapitulated subtle differences in waveforms between various cardiac diseases, which could provide a powerful platform to study the mechanical consequences of cardiovascular disease and cardiac drugs on cardiac contraction and endothelial cells. Follow-up studies of molecular and genetic features of cells within the Adapt-Pump system could improve our understanding of cardiac and endothelial cells under physiological and pathological conditions, guiding improved treatments for patients.

## Materials and methods

### hPSC culture and cardiac progenitor differentiation

Human LQTS iPSCs (iPSCORE_2_1^44^) and human ESCs (H1 and H9^45^) were maintained on Matrigel (Corning)-coated surfaces in mTeSR1 (STEMCELL Technologies) as previously decribed^46^. Cardiac progenitor differentiation was performed as described previously^22^. Briefly, hPSCs were singularized with Accutase (Thermo Fisher Scientific) and plated onto Matrigel-coated plates at a density of 3.0×10^5^ cells/cm^2^ in mTeSR1 supplemented with 10 μM Rho-associated protein kinase (ROCK) inhibitor Y-27632 (Selleckchem) on day −2. The differentiation was initiated with Wnt activation by 8 μM CHIR99021 (Selleckchem) on day 0 and sequential Wnt inhibition by 5μM IPW2 on day 3. Cardiac progenitors were ready on day 6 for CS generation.

### Cardiac spheroid generation for imaging

Cardiac progenitors at differentiation day 6 were singularized with Accutase. 3000 cardiac progenitors were resuspended in 30 μL cell culture medium with 2.4 mg/mL methylcellulose (Sigma Aldrich) for each CS. The mixed resuspension was then plated on a 60 mm tissue culture dish as a droplet and the tissue culture dish was flipped for the suspension to form an spheroid. After 48 hr, the individual spheroid were collected with a 200 μL pipette tip and embedded into Matrigel. The cardiac contraction imaging was performed on day 30. For drug treatment studies, individual spheroids were imaged over 60 seconds, and then spheroids were dosed with either 1 μM nifedipine (Sigma Aldrich) or 1 μM verapamil (Sigma Aldrich) for 10 minutes before imaging again.

### EC progenitor differentiation and EC long-term shear stress treatment

EC progenitors were differentiated as described previously^34^. Briefly, IMR90-4 hPSCs were singularized with Accutase and seeded onto a Matrigel-coated plate at a density of 2×10^4^ cells/cm^2^ in mTeSR1 supplemented with 10 μM ROCK inhibitor Y-27632 on day −3. Differentiation was initiated by Wnt signaling activation with 8 μM CHI99021 for 48 hours in LaSR basal medium. After 2 days, cells were maintained in LaSR basal medium for an additional 3 days. On day 5, CD34+ EC progenitors were purified with an EasySep Magnet kit (STEMCELL Technologies) with a CD34 antibody. Purified CD34+ EC progenitors were then seeded onto a 0.2 mm ibidi μ-slides or an ibidi 24-well μ-plate precoated with 10 μg/mL Collagen IV (Sigma Aldrich). 24 hours after the cells were seeded, pulsatile shear stress was applied to the cells for 48 hours.

### Immunochemistry

Cells were rinsed with ice-cold PBS (Thermo Fisher Scientific) twice and fixed in 4% paraformaldehyde (PFA) for 15 minutes. Cells were then blocked and permeabilized with 10% goat serum in PBS containing 0.3% Triton X-100 for 30 minutes (10% PBSGT). Primary antibodies (CD31, 1:100, Invitrogen, 14-0319-82, PIEZO1, 1:100, Invitrogen, MA5-32876) were diluted in 10% PBSGT, and cells were incubated in the diluted primary antibodies overnight at 4°C. Then cells were rinsed with room temperature PBS three times and incubated with secondary antibodies (Secondary antibodies, 1:200, Goat anti-Mouse IgG2a Alexa Fluor 488, Goat anti-Mouse IgG1 Alexa Fluor 594) in 10% PBSGT at room temperature for 1 hour. After incubation, cells were washed with PBS three times followed by treatment with DAPI Fluoromount-G (Southern Biotech). Cells were imaged with a Leica SP8 3×STED superresolution microscope.

### Flow cytometry

Cells were disassociated with Accutase, fixed in 1% PFA for 15 minutes at room temperature, and then blocked with 0.5% bovine serum albumin (BSA) with 0.1% Triton X-100 for non-permeabilization or without 0.1% Triton X-100 for membrane permeabilization. Cells were then stained with conjugated CD31 (Miltenyi Biotec, 130-119-891) in 0.5% BSA with or without 0. 1% Triton X-100. Data were collected on a FACSCalibur flow cytometer.

### Extraction of contraction waveforms

Quantitative brightfield imaging of CSs was performed using a Nikon Ti-U with a 4× air objective (0.13NA). A high-speed monochrome camera (Basler ace acA1300-200 μm, 1280×1024 pixels) captured image time-series of CS contractions over 60 seconds at 30 fps. A custom algorithm was used to analyze these image time-series and extract the pulsatile waveform from the CS contractions. The algorithm first segments the CS in every frame to generate a mask to measure the spheroid area. An initial CS mask was generated using graph-cut segmentation *(lazysnapping,* MATLAB) of an image calculated from the average of all frames. Mathematical morphology operations were used to refine the mask and generate a single segmented region containing no holes. This initial mask was then applied to each frame to exclude regions outside the fully relaxed spheroid. This enabled intensity-based segmentation *(multithresh, imquantize,* MATLAB) of the spheroid in each frame of the image time-series to measure the area of the CS *(regionprops,* MATLAB). The extracted area waveform captures the contraction characteristics of each CS at high temporal resolution and minimal computational cost. All analysis was performed in MATLAB 2018b (MathWorks).

### Generation of pressure waveform

A pressure waveform was generated from the contraction waveform via the following equation: *Pressure* = *C* * (*Area_i_* – *Area_reiaxed_*), where *C* is a scaling factor (units: mbar/mm^2^), *Area_i_* is the area of the spheroid mask at each time point, and *Areareiaxed* is the area of the mask when the spheroid is fully relaxed. For the pulsatile flow characterization, a peak pressure of 150 mbar was set for all waveforms. For EC differentiation, a scaling factor of *C* was calculated that converted one of the contraction waveforms into a pressure waveform (range: 0 to 800 mbar). The scaling factors for all other waveforms were calculated so that the average pressure of all waveforms was the same.

### Pump hardware

Pulsatile and steady-state flows were generated using a pressure controller (Elveflow OB1 MK3) connected to building vacuum (−1930 mbar) and pressurized air (6890 mbar). This controller provided the fast pressure modulation (9 ms settling time, 35 ms response time, 122 μbar pressure resolution) required to recapitulate the spheroid contraction waveforms. The controller used in this study was configured with two channels with output pressures from −900 mbar to 1000 mbar. For the flow validation studies, one of these channels was connected to a reservoir and used to generate unidirectional flow through the flow channel. For long-term experiments, a recirculation circuit was used that comprised two channels each connected to a reservoir and a flow switch (Elveflow MUX Injector), which is a 6-port/2-position valve that allows rapid switching between the two configurations. This allowed for unidirectional flow to be maintained through the flow channel while the medium recirculated between the two reservoirs. Teflon PTFE tubing (1/16” inner diameter) was used to carry medium between the reservoirs and the flow channel. A custom script was used to control the pressure controller and flow switch to generate pulsatile flow through the flow channel. All pump control was achieved using MATLAB 2018b (MathWorks).

### Particle flow validation and streak analysis

1 mg/mL of 4 μm fluorescent microbeads in DMEM phenol red-free medium in a 0.8 mm-ibidi μ-slide were flowed at constant flow rates generated by applied pressures of 50 mbar, 100 mbar, 150 mbar, and 200 mbar. Bead flowing images were collected with a Nikon Ti-U epifluorescence microscope with a Lumencor SOLA FISH light source (Ex/Em 480/520nm). Images were collected at the z-positions of defined by the 10× (0.25 NA) Nikon objective position at 0.5 mm, 0.6 mm, 0.7 mm, and 0.8 mm, where position 0 mm is the bottom surface of the channel. Images were collected at 200 fps using a scientific CMOS camera (Flash 4, Hamamatsu). To achieve 200 fps acquisition, 4× on-camera pixel binning and a cropped region-of-interest reduced the size of the acquired image to 256 pixels × 128 pixels. This rectangular region-of-interest was oriented in the center of the channel with the bead flow parallel to the long-axis (256 pixels).

A custom algorithm was developed to measure the speed of fluorescent beads moving through the channel. The algorithm is based on particle streak velocimetry, which analyzes the velocity of beads moving faster than the frame period by measuring the length of particle streaks. This enables the quantification of >1cm/s flow using widely available CMOS cameras. To extract the speed at which the beads were moving through the microfluidic channel in each frame of image times-series, elongated or tubular structures were enhanced using Hessian-based multiscale filtering *(fibermetric,* MATLAB). A threshold was calculated based on Otsu’s method and applied to the Hessian-enhanced image time-series *(graythresh, imquantize,* MATLAB), which successfully segmented each streak to generate candidate streak masks. These candidate streak masks were then filtered based on their morphology, and small objects and streaks touching the image border were removed *(regionprops, imclearborder,* MATLAB). The lengths of the remaining streaks were measured in units of micrometers. The speed of each streak was calculated by dividing the length of each streak by the frame period (5ms). The pressure waveform and its corresponding bead velocity waveform were synchronized by finding the shift value that maximized the crosscorrelation between the two waveforms *(finddelay, alignsignal,* MATLAB). All analysis was performed using MATLAB 2018b (MathWorks).

### Computational fluid dynamics simulation

Autodesk Inventor was used to draw the microfluidic channel for the microfluidic channel simulation. The shape was then imported into Autodesk CFD 2019 for the simulation. The channel dimensions for the bead analysis were defined as 50×0.5×0.8 mm (Length×Width×Height, ibidi μ-slide used for bead speed validation) and 50×0.5×0.2 mm (Length×Width×Height, ibidi μ-slide used for long-term cell culture), respectively. The fluid viscosity was set to 0.00072 Pas, which is the value for standard cell culture medium at 37°C (ibidi pump manual). The environment temperature was set to 37°C. The boundary condition values for the simulation were set to the conditions previously calculated in the bead experiment to calculate shear stress (parameters: inlet and outlet flow rate 40 mL/min, inlet gauge pressure 50 mbar, outlet pressure 0 mbar).

To analyze the results, three planes were added to the simulation: speed (cm/s), shear stress (dyne/cm^2^), and pressure (mbar). To simulate the laminar flow section of the channel, these planes were placed near the center length of the channel simulation (z-axis) at 25 mm from the channel inlet and outlet. Values from five places within the channel were also verified: center of channel (0, 0, 25 mm), bottom center of channel (0, −0.4, 25 mm), left center of channel (2.5, 0, 25 mm), top center of channel (0, 0.4, 25 mm), and right center of channel (2.5, 0, 25 mm).

### Real-time rtAdapt-Pump implementation

CSs were embedded in Matrigel and plated into the microchannel of a 0.8 mm ibidi sticky μ-slide (lower chamber). The sticky slide was then sealed with polyester membrane and covered with another 0.8 mm ibidi sticky μ-slide (upper chamber). The upper chamber was filled with CS culture medium. The CS was imaged with a Nikon Ti-U with a 10× (0.25 NA) objective at 20 fps under static condition over 30 seconds to measure the contraction waveform at baseline. Then, the CS was imaged for an additional 30 seconds with pulsatile flow generated in real-time based on the pulsatile contraction waveforms also extracted instantaneously. This workflow was achieved through integrating quantitative imaging of CS, extraction of pulsatile contraction waveform, generation of pressure waveform, and pump control into a single real-time framework. Each part was designed to work with low computational cost so that the real-time framework operated at 20 frames per seconds. Comparison of the contraction waveform at baseline and with real-time flow stimulation was achieved through a custom script written to quantify the peak amplitude, contraction duration, and beat frequency. This real-time framework was realized using MATLAB 2018b (MathWorks).

### Statistics

Data are presented as means ± SEM. Statistical significance was determined by Student’s t test (two-tailed) between two groups. Three or more groups were analyzed by one-way analysis of variance (ANOVA) followed by Tukey’s post hoc tests. P < 0.05 was considered statistically significant and indicated in the figures.

---

**Figure S1.**
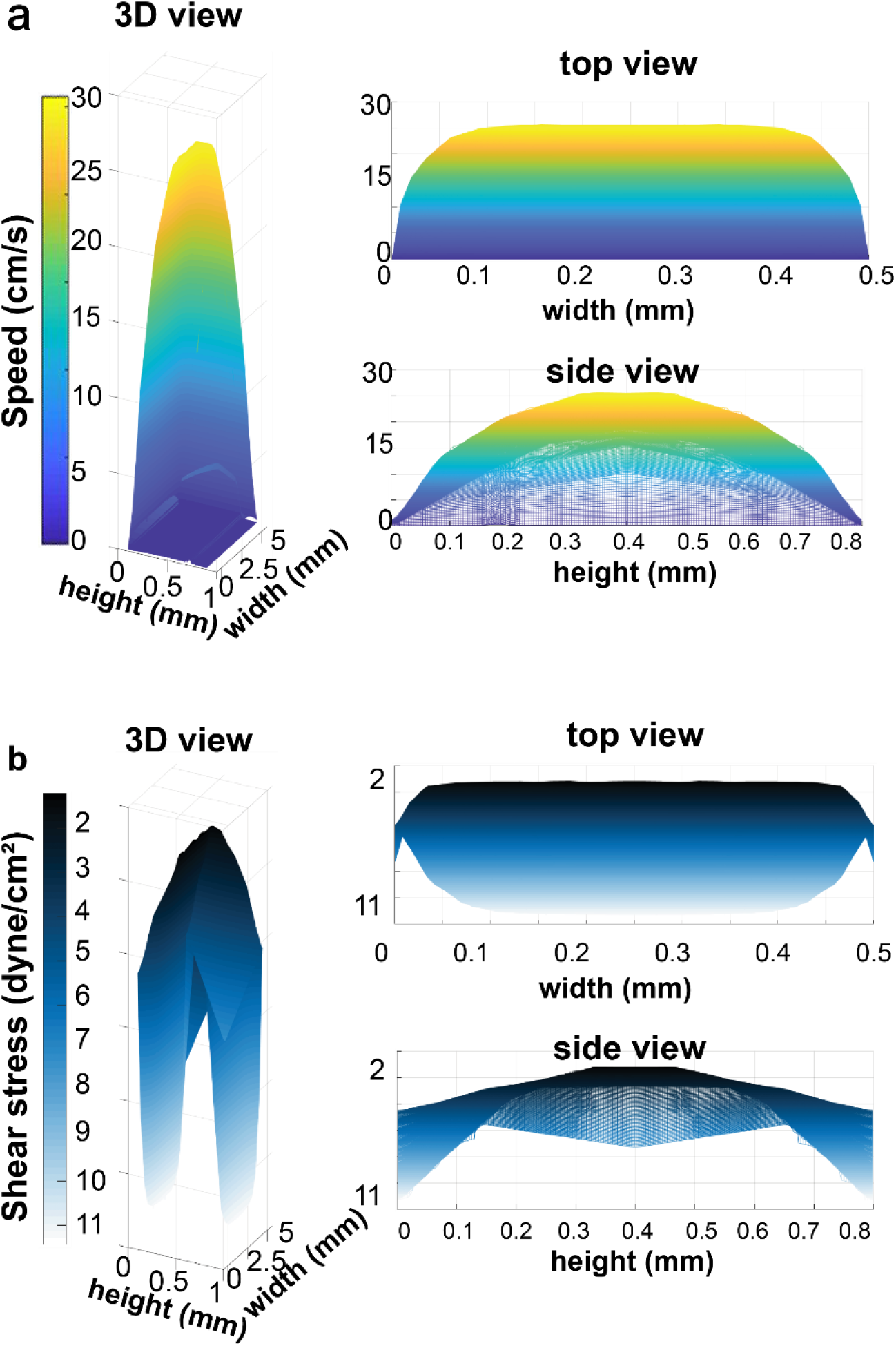
Shear stress and speed simulations. 3D plot of shear stress and speed simulations at 800 mbar pressure using the dimensions of a 0.8 mm ibidi μ-slide.

**Figure S2.**
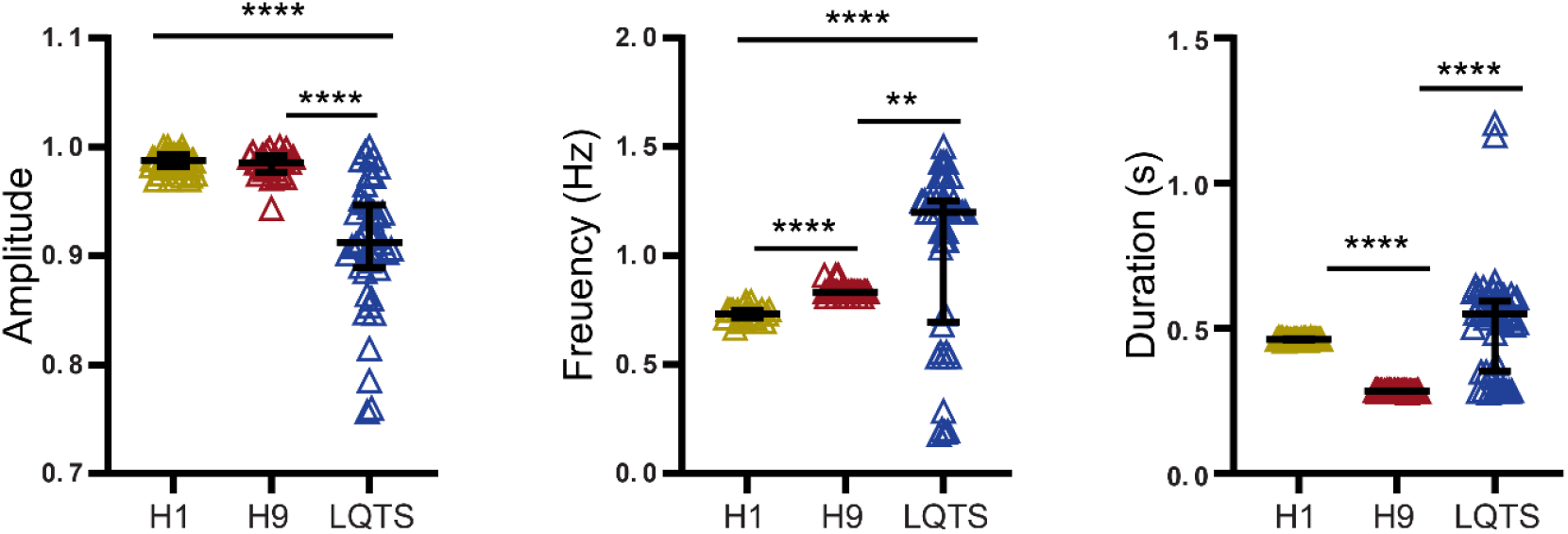
Characterization of different hPSC-CS contraction waveforms. Normalized contraction amplitude, contraction frequency, and contraction duration are presented as mean ± SEM. Each hPSC-CS contraction waveform was calculated from the individual contractions over 60 seconds. Statistical significance was determined by one-way analysis of variance (ANOVA) followed by Tukey’s post hoc tests. \*\**p*< 0.01, \*\*\*\**p* < 0.0001.

**Figure S3.**
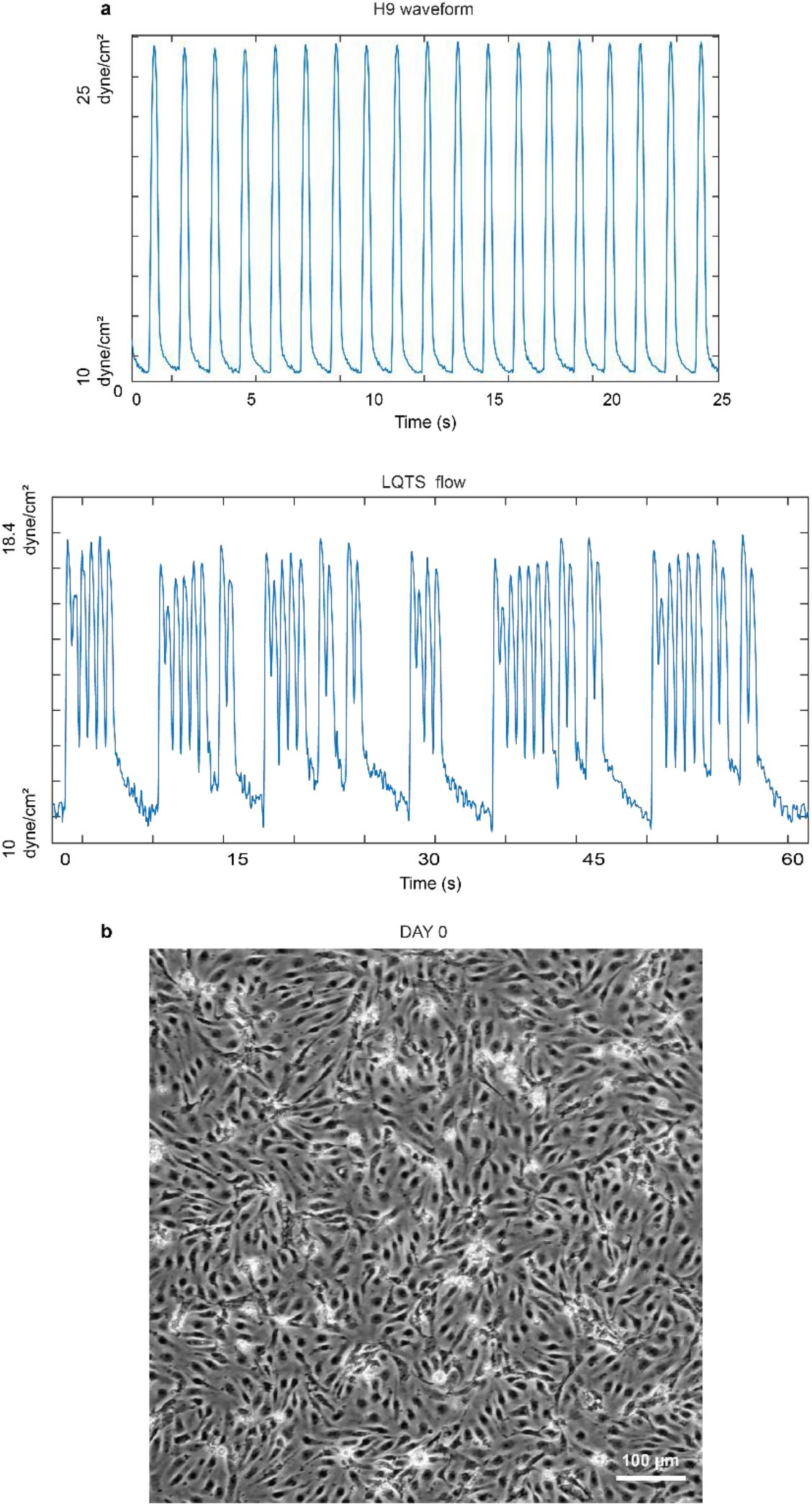
Pulsatile flow profile applied to EC differentiation and brightfield images of EC after differentiation under shear stress. (**a**) Pulsatile flows applied to EC progenitors during differentiation. H9-CS derived pulsatile flow and LQTS-CS derived pulsatile flow applied to the EC progenitors were maintained at the baseline of 10 dyne/cm^2^. The peak shear stress for H9-CS derived pulsatile flow was 25 dyne/cm^2^ and the peak shear stress for LQTS-CS derived pulsatile flow was18.4 dyne/cm^2^. The average shear stress levels for both hPSC-CS derived pulsatile flows were at 13.7 dyne/cm^2^. (**b**) Representative brightfield images of EC progenitors (DAY 0) Scale bar, 100 μm.

**Supplemental Table 1.** Pulsatile waveforms and their corresponding contraction waveforms are highly correlated. Pearson correlation was performed to verify the cross-correlation between pulsatile waveforms and their corresponding contraction waveforms. R values indicated.

| Group | Pearson's r value |
| --- | --- |
| H1 ES-CS | 0.96 |
| H9 ES-CS | 0.96 |
| LQTS iPSC-CS | 0.98 |
| LQTS-CS before nifedipine | 0.79 |
| LQTS-CS after nifedipine | 0.83 |
| H9-CS before verapamil | 0.97 |
| H9-CS after verapamil | 0.95 |

Supplemental videos: 21 supplemental videos

## Acknowledgements

### Funding

This work was supported by the Morgridge Institute for Research and the NSF Center for Cell Manufacturing Technologies (grant EEC-1648035).

### Author contributions

The project was conceived by T.Q., and M.C.S. The Adapt-Pump platform was designed and implemented by D.A.G. and T.Q., and image analysis algorithms were developed by D.A.G. and K.E.T. The experiments were designed by T.Q., D.A.G., and M.C.S. and were carried out by T.Q., D.A.G., E.C.G., B.D.G., and K.E.T. The manuscript was written by T.Q., D.A.G, S.P.P., and M.C.S. We thank assistance from E.C.G., B.D.G., and K.E.T. on this project.

### Competing interests

All other authors declare that they have no competing interests.

### Data and materials availability

All data needed to evaluate the conclusions in the paper are present in the paper and/or the Supplementary Materials. Additional data related to this paper may be requested from the authors.

